# A structural analysis of amyloid polymorphism in disease: clues for selective vulnerability?

**DOI:** 10.1101/2021.03.01.433317

**Authors:** Rob van der Kant, Nikolaos Louros, Joost Schymkowitz, Frederic Rousseau

**Affiliations:** Switch laboratory, VIB Center for Brain and Disease Research, Leuven, Belgium; Department for Cellular and Molecular Medicine, KU Leuven, Herestraat 49, 3000 Leuven, Belgium

## Abstract

The increasing amount of amyloid structures offers an opportunity to investigate the general principles determining amyloid stability and polymorphism in disease. We find that amyloid stability is dominated by about 30% of residues localized in few segments interspersed with regions that are often structurally frustrated in the cross-β conformation. These stable segments correspond to known aggregation-nucleating regions and constitute a cross-β structural framework that is shared among polymorphs. Alternative tertiary packing of these segments within the protofibril results in conformationally different but energetically similar polymorphs. This combination of a conserved structural framework along the axis and energetic ambiguity across the axis results in polymorphic plasticity that explains a number of fundamental amyloid properties, including fibril defects and brittleness but also the polymorphic instability of amyloids in simple aqueous buffers. Together these findings suggest a structural model for in vivo polymorphic bias and selective cellular vulnerability whereby (1) polymorphic bias is induced by particular templating interactions in susceptible cells, (2) once formed specific polymorphs are entropically primed to selectively bind similar targets in neighbouring cells, (3) conservation of polymorphic bias during pathological spreading implies the continued presence of similar templating interactions in successive susceptible cells and (4) absence of templating interactions relaxes polymorphic bias possibly allowing for the modification of cellular susceptibilities during disease progression by novel templating interactions.

## Introduction

Amyloids are the main constituent of pathological protein deposits in more than thirty diseases including capital neurodegenerative disorders (e.g. Alzheimer’s Disease) and other major pathologies (e.g. type II diabetes) or systemic forms such as light-chain amyloidosis^1^. These proteins are completely dissimilar in terms of sequence, native structure, and function. However, their amyloid conformation shares a common structural basis, the so-called cross-β fibrillar architecture^2^. Distinct amyloid pathologies are associated to the deposition of particular proteins in specific disease-vulnerable cells and tissues^3,4^. Even more remarkably, amyloid conformational variants of the same protein called polymorphs seem to be associated to distinct pathologies, e.g. tauopathies, including diseases such as Alzheimer disease, Chronic Traumatic Encephalopathy, Progressive Supranuclear Palsy or Pick’s disease which originate in separate areas of the brain and are associated to different structural polymorphisms of the protein tau^5^. Although the general properties of amyloid structure are well established, our structural understanding remains insufficient to mechanistically connect their structural features to molecular mechanisms of disease including the origin of amyloid toxicity and the role of amyloid polymorphism in selective cellular vulnerability.

Recent developments in both Solid State NMR (ssNMR) and cryo-electron microscopy (cryoEM) have resulted in an explosion of amyloid structures from both *in vitro*, as well as *in vivo* assembled amyloid fibrils. The amount of structures now available is reaching what it was at the beginning of the 1980s for globular proteins (around 100 structures) which was sufficient to shape the first insights on general principles determining globular architecture and its more fine-graded implications for protein function and evolution (**Fig. 1A-1O**). How well do disease-associated amyloid structures explain some of their defining features, such as their high thermodynamic stability, their polymorphic nature but also the sensitivity of polymorphism to environmental conditions? Can these structures also provide insights towards the origin of fibrillar growth defects and fiber brittleness, the tendency of amyloids to populate oligomeric intermediates, the affinity for metals and the water activity of amyloids? Can such structural insights, in turn, help design experiments and guide our search for the molecular mechanisms contributing to amyloid toxicity and the role of amyloid polymorphism in the process of selective cellular vulnerability?

**Figure 1.**
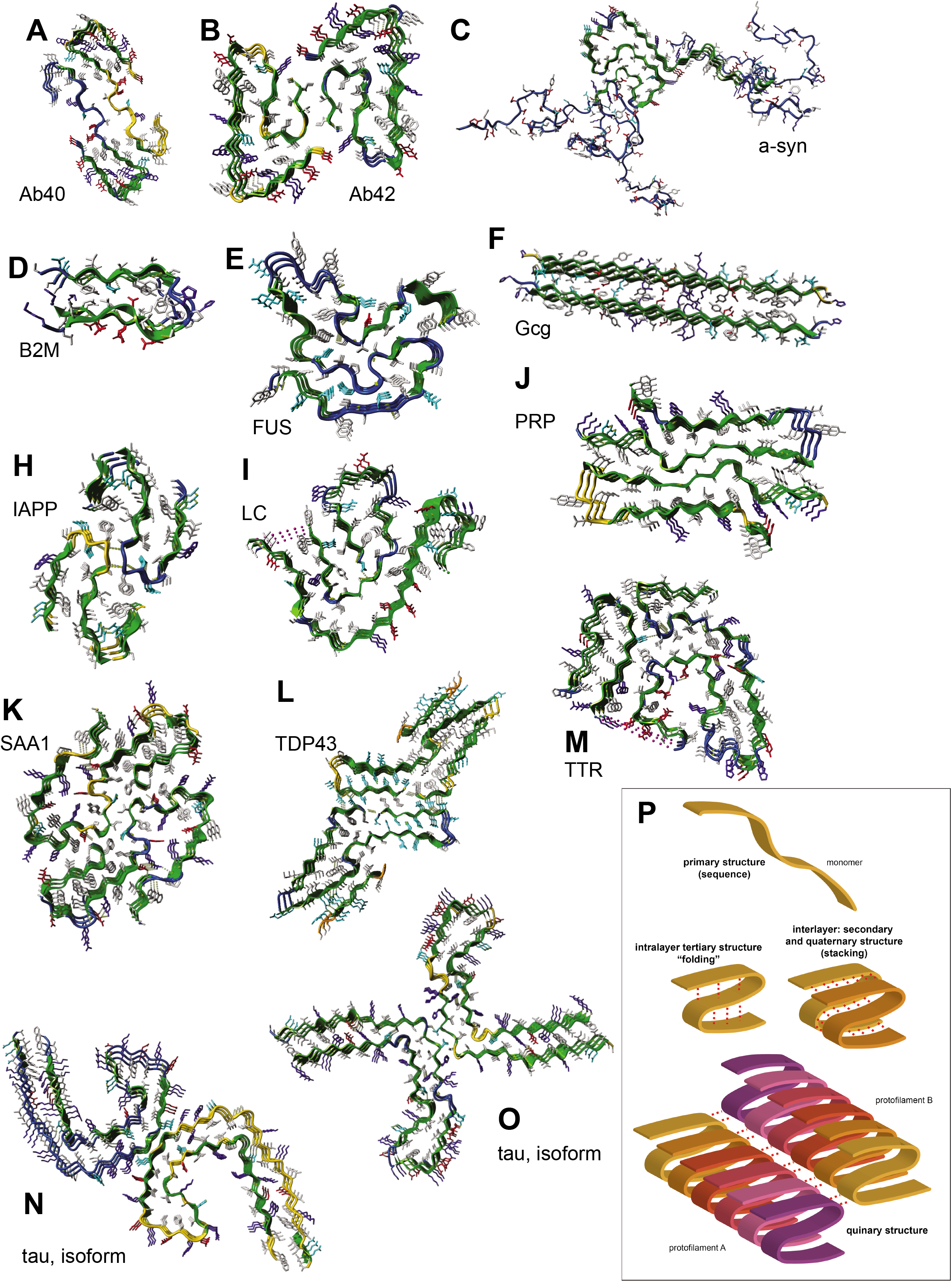
Defining the structural hierarchy of amyloid fibril structures. (**A-O**) Representative structural layouts of fibrils formed by several amyloid-forming proteins that were included in our study (**Supplementary Table 1**). (**P**) Complexity of the different folding levels of amyloid fibrils. Protofibrilar assembly is defined by a series of intralayer packing interactions that hold together the tertiary fold with secondary backbone intermolecular interaction forming their quaternary structure. Protofibrillar surface interactions introduce a quinary layer of complexity that contributes to amyloid polymorphism.

Here, we performed a global structural analysis of 66 disease-associated amyloid structures and compared structural and thermodynamic characteristics between amyloids but also between amyloids and globular proteins. Using FoldX^6^, we further performed a more in depth *in silico* thermodynamic analysis of the Amyloid-beta (Aβ peptide, tau and alpha-synuclein (α-syn) for which polymorphic structural information is available including *ex vivo* patient amyloid structures. Our findings highlight how the conjunction of common motifs of quaternary structural stability along the fibril axis and tertiary structural ambiguity across the fibril axis explain both the stable and robust propagative nature of amyloids, as well as the high sensitivity of polymorphism to environmental conditions. It also helps explaining the polymorphic instability and the intrafibrillar polymorphic transitions observed in simple aqueous buffers^7^. We discuss how these findings suggest that the maintenance of a more robust polymorphic bias observed in specific pathologies must result from the presence of consistent polymorph-selecting interactions in successive vulnerable cells. Finally, we also discuss how both metals and structural waters help accommodate unfavourable charged side chain stacking, as well as unsatisfied main-chain H-bonds. These interactions could potentially modify both metal homeostasis, as well as water activity in disease.

## Results

### Protein Structural Hierarchy in Amyloid Structure

Amyloid structure displays very different properties along versus across the fibril axis. As a result, the protein structural hierarchy commonly used to describe globular protein structure^8,9^cannot be transferred to amyloid structure^8,9^. First, ‘classic’ secondary structure does not exist in the amyloid state. Instead, main-chain H-bond saturation is here provided by quaternary H-bonds that staple adjacent chains into the archetypal amyloid cross-β sheet running perpendicular to the fibril axis. Second, while short amyloid peptides can organise into both parallel and antiparallel β-sheets^10,11^, longer or full-length protein fibrils are so far primarily found forming parallel β-sheet structures. As a result, side chain interactions along the fibre axis result mainly from the registered stacking of identical side chains, a situation that is unique to amyloid assembly. Third, amyloid tertiary structure results from invagination of the cross-β sheet, whereby distinct segments of the cross-β sheet pack against each other to form steric zippers. Rungs of the amyloid fibril can run strictly perpendicular to the fibril axis so that tertiary interactions are mainly formed between sidechains within the same layer. Alternatively, rungs can be slanted so that each monomer interacts with several subsequent layers. Contrary to the strict in-register quaternary structure of the cross-β sheet, tertiary amyloid structure is more versatile and a source of polymorphism, as alternative tertiary folds can be observed for a single amyloid sequence. The final topology of a single monomer within the tertiary fold often adopts a serpentine shape at first sight, reminiscent of the 2D projections of the Greek key motifs that are found in globular proteins. However, in globular structures, Greek key topology is defined by its intramolecular main chain-main chain H-bond network between β−sheets. Conversely serpentine-like topology in amyloids is defined by its tertiary sidechain contact map. A final distinction between globular and amyloid structural hierarchy is the fact that individual amyloid protofibrils can co-assemble to form amyloid fibrils that are stabilized by non-covalent interactions between constituting protofibrils. Alternative fibrillar assemblies between identical protofilaments constitute another source of amyloid polymorphism (**Fig. 1P, quinary structure**).

### The stability of amyloids is determined by a minority of residues that favour cross-β structure

Overall, cryoEM structures have Ramachandran maps that are very similar to those observed in high resolution X-ray structures (**Fig. 2A and B**). Solid state NMR structures, however, often adopt improbable backbone dihedral angles (**Fig. 2C**). As inaccuracy in backbone conformation precludes reliable energy estimation with FoldX, we separately considered ssNMR structures (**Supplementary Table 1, sheet ‘Fibril structures’**). In order to evaluate the proportion of residues that contribute to the stability of amyloid structures and compare it with globular structure, we calculated the per residue distribution of free energy contributions of all residues in both datasets. For globular structures (**Fig. 2D**, blue) we find a biphasic distribution, with roughly one third of the residues contributing strongly to the stabilisation of these structures (negative ΔG), with the remaining residues being neutral or slightly destabilising. As can be expected, the strongly stabilising residues are part of the hydrophobic core (**Fig. 2E**) and mostly reside in secondary structure elements (**Fig. 2F**) that are satisfied by mainchain H-bonds (**Fig. 2G**). Neutral or destabilizing residues, on the other hand, are mostly found in loops that are more exposed (**Fig. 2E**), possess unsatisfied backbone H-bonds (**Fig. 2G**) and are sometimes structurally frustrated, as they more frequently adopt less favourable dihedral angles (**Fig. 2H**).

**Figure 2.**
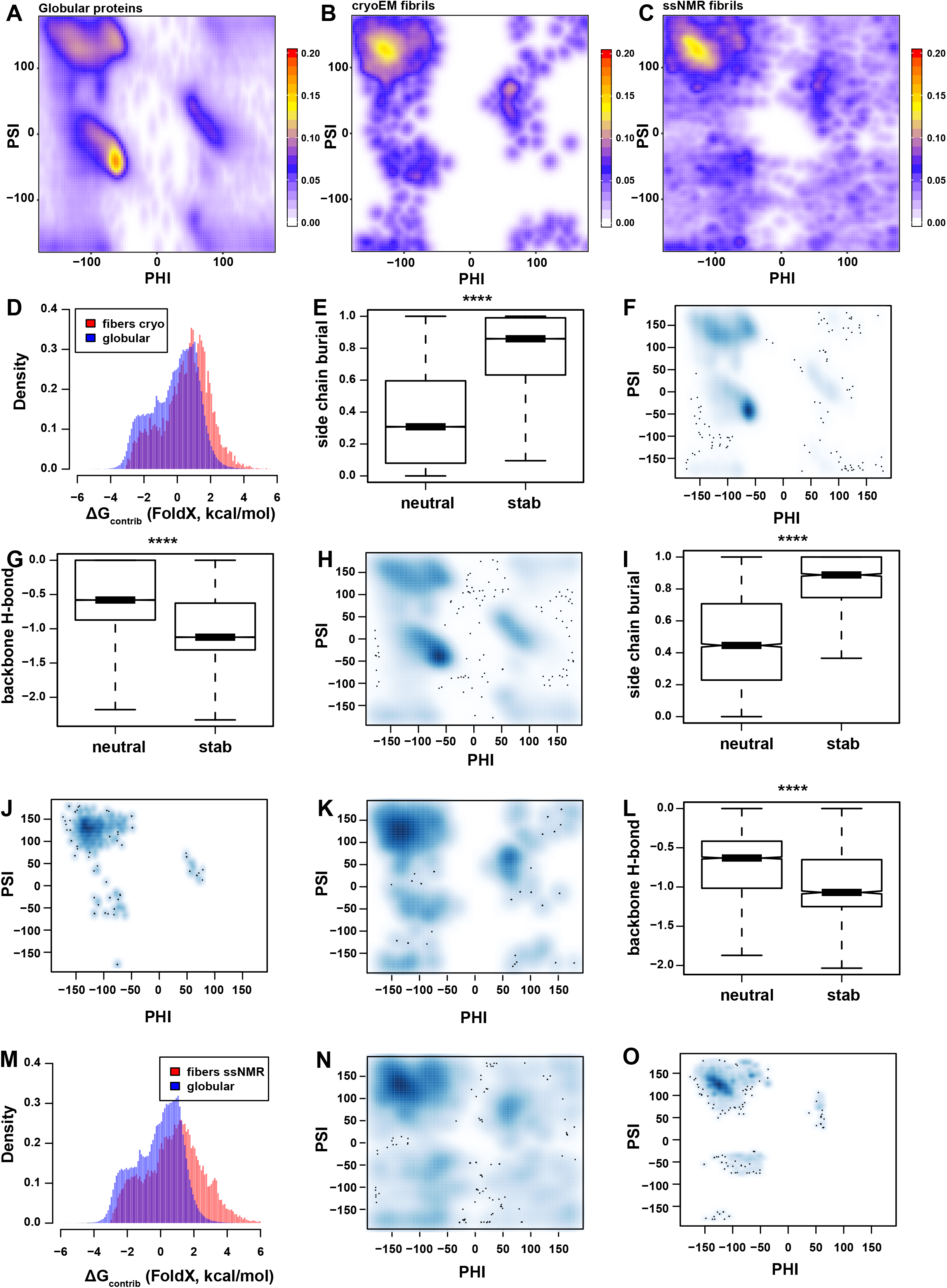
Structural assessment and comparison of amyloid fibril structures to globular folds. **(A-C**) Ramachandran density plots indicating regions of dihedral angles occupied by residues in the reference set of globular protein folds (**A**), compared to residues in (**B**) cryoEM and (**C**) ssNMR structures of amyloid fibrils. (**D**) Distribution analysis of per residue energy contributions in globular folds (shown in blue) and cryoEM fibril structures (shown in blue). (**E-H**) Side chain burial (**E**), backbone hydrogen bond energies (**G**) and Ramachandran plot comparisons for stabilising (**F**) and neutral or slightly destabilising residues (**H**) of globular folds. (**I-L**) Side chain burial (**I**), backbone hydrogen bond energies (**L**) and Ramachandran plot comparisons for stabilising residues (**J**) and neutral or slightly destabilising (**K**) of cryoEM amyloid fibril structures. (**M-O**) Distribution analysis of per residue energy contributions in globular folds (shown in blue) and ssNMR fibril structures (shown in blue). Ramachandran density plots for neutral or slightly destabilising (**N**) and stabilising residues (**O**) of cryoEM amyloid fibril structures. Significance levels for boxplots are indicated (* for P ≤ 0.05, ** for P ≤ 0.01, *** for P ≤ 0.001, **** for P ≤ 0.0001).

We find that fibril structures possess a very similar biphasic distribution of residue free energy contributions as globular structures, with again about one third of residues favourably contributing to the stability of amyloid structures (**Fig. 2D**, red). The slight deviation towards poorer energies is likely a reflection of the lower overall resolution of these structures, compared to the reference set of globular protein structures (**Supplementary Table 1, sheet ‘Globular structures X-ray’**). Amyloid stabilizing residues are more buried than neutral and destabilizing residues (**Fig. 2I**) and exclusively found in theβ-sheet region of the Ramachandran map, as expected for the cross-β amyloid structure (**Fig. 2J**). Neutral and destabilizing residues, on the other hand, show sometimes pronounced deviation from optimal β-sheet dihedral angles (**Fig. 2K**) and also display less backbone hydrogen bonding (**Fig. 2L**). A similar distribution of residue free energy contributions was observed for the ssNMR structures (**Fig. 2M**), with the tail of poor energies being even more populated, likely due to the slightly lower structural quality also observed in the Ramachandran map (**Fig. 2C**). However, despite their lower resolution, the stabilising residues in NMR structures still populate a very narrow area of the β-sheet region of the Ramachandran map (**Fig. 2N & O**). Together, this analysis shows that on average only about one third of the residues in amyloids are responsible for the thermodynamic stability of these structures and that most residues are either neutral or even thermodynamically unfavourable, proportionally similar to those observed in globular structures.

### Amyloids possess segments of high structural stability and regions of significant structural frustration

Amyloid protofibrils are constituted of quaternary cross-β sheet of aligned monomers with stacked identical side chain residues along the fibril axis. However, the tertiary fold of amyloid structures also contains turns and often cavities. As a result, not all regions of the amyloid can be expected to equally contribute to the stability of amyloid fibrils. Overlap of the energy profiles of individual polymorphs revealed that they are very similar to each other with the same sections of the protein sequence constantly determining the stability of the different polymorphs (**Fig. 3A, Supplementary Fig. S1 and S2**). To quantify this, we calculated the pairwise correlation matrix between the energy profiles of each polymorph (**Fig. 3B**), revealing median correlation values between the polymorphs of 0.63 for α-syn, 0.69 for tau and Aβ42 and 0.95 for Aβ40, which formally confirms their similarity. The correlation in α-syn or tau is somewhat lower than in shorter amyloids, such as Aβ, due to differences in length of these amyloids.

**Figure 3.**
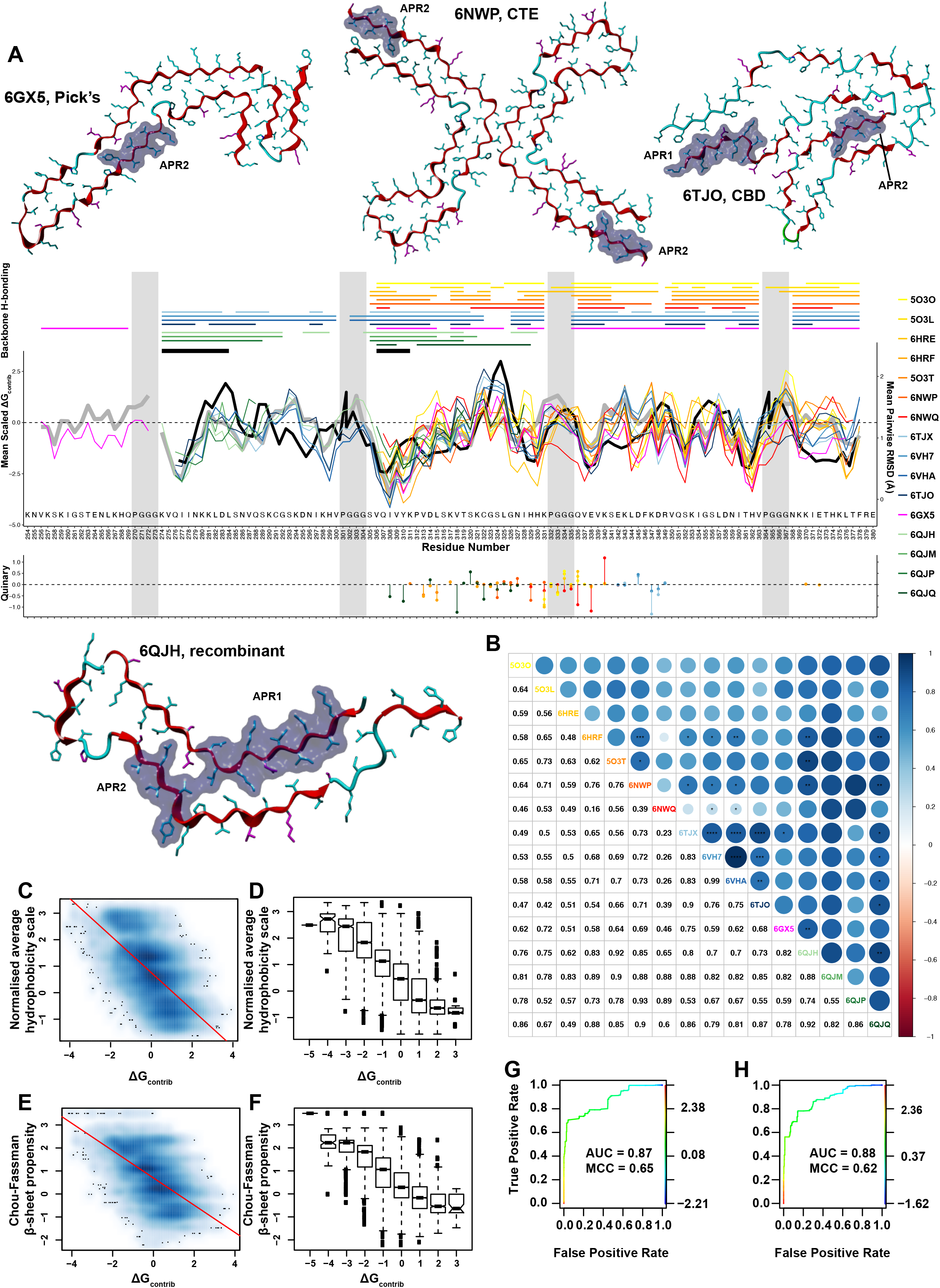
Cross-comparison of defining features between amyloid structural polymorphs. (**A**) Representative structures of tau polymorphs, with positions of known APR segments highlighted as black segments. Distributions of stability free energy contributions (middle plot) are aligned to quinary energy contributions (bottom plot) and overlaid in different colours for different structural polymorphs of tau. The top plot indicates hydrogen bond patterning (colour-coded per polymorph) and the position of experimentally determined APRs (black segments). RMSD distribution, produced after superimposing all structures, is plotted as a thick black line in the middle plot. Shaded boxes indicate turns that connect tau repetitive segments. (**B**) Pairwise comparison plot of individual tau polymorphs. Top-half indicates strong structural similarities by circle sizes (colour-coded from red to blue) and bottom half indicates pairwise scores. (**C-D**) Per residue density distribution (**C**) and binned box-plot analysis (**D**) of hydrophobicity, plotted against free energy contributions (red line indicates the linear fitting). (**E-F**) Per residue density distribution (**E**) and binned box-plot analysis (**F**) of β-sheet propensity, plotted against free energy contributions (red line indicates the linear fitting). (**G-H**) ROC analysis of (**G**) β-sheet propensity and (**H**) hydrophobicity for the prediction of APRs in the amyloid core of structural polymorphs.

Secondly, we find that the most stable structural elements are also those displaying the lowest backbone RMSD variation between polymorphs (**Fig. 3A, Supplementary Fig. S2,** black lines), showing that similar energetic profiles indeed result from identical stabilizing structural elements. To probe the determinants of these common stable segments further, we analysed the relationship between hydrophobicity^12^in windows of five amino acids and the contribution of those amino acids to the fibril structure (**Fig. 3C**), revealing a clear correlation (adjusted R-squared = 0.40, p < 2.2e^-16^). Binning of the ΔG values and averaging the propensity scores over these bins showed that the relationship is sigmoidal in nature (**Fig. 3D**) with saturation of the hydrophobicity occurring at both ends of the ΔG_contrib_ spectrum. The other well-known determinant of amyloid formation is local β-sheet propensity (Chou-Fassman parameter^13^). Indeed, a similar relationship can be found as for hydrophobicity (**Fig. 3E**, adjusted R-squared = 0.35, p < 2.2e^-16^) and after binning this relationship equally appears sigmoidal (**Fig. 3F**). This shows that the structural stability is highest in the sequence segments that have the highest propensity of β-sheet conformation, and that these regions form the most regular β-sheet structure in the mature fibrils (**Fig. 2O**). In fact, a simple linear model using only these two parameters can predict the ΔG_contrib_ values in these fibrillar structures with a correlation of 0.59 (p < 2.2e^-16^). This shows that the stability of the fibrils is determined in a similar manner to globular proteins, namely by burying hydrophobic groups with a high propensity for mainchain H-bond saturation. But, as in globular structures, this is not possible for every part of the sequence. As a result, only about one third of the sequence contributes to the stability of amyloid fibrils while the other two thirds are either neutral or even energetically unfavourable for the amyloid structure (**Fig. 2D**).

Thirdly, we observe a remarkable overlap between the segments determining the stability of amyloid structures with those determining the kinetics of amyloid assembly. We annotated the energy profiles with the aggregation-prone regions (APRs) that were experimentally determined to be essential for the kinetics of amyloid formation by their ability to either form amyloid structures in isolation or by their ability to specifically seed amyloid aggregation of full length peptides and proteins^11,14^(**Fig. 3A, Supplementary Fig. S1 and S2,** black segments). To quantify the impact of β-sheet propensity and hydrophobicity in defining the APRs, we computed Receiver-Operator Curves (ROCs). The analysis shows that the experimentally validated APRs can indeed be identified within these sequences with accuracy close to current state-of-art predictors^15^ from either β-sheet propensity (**Fig. 3G**, AUC = 0.87, MCC = 0.65) or hydrophobicity (**Fig. 3H**, AUC = 0.88, MCC = 0.62). Together, these findings demonstrate that despite their polymorphism amyloid structures of a specific protein sequence are organised around a common framework of segments that favour stable cross-β structure along the fibril axis and that these elements are the same that are required for amyloid nucleation and elongation.

### Polymorphism results from degenerate tertiary packing against a common stable cross-β framework

Most structural information on amyloid polymorphism is available for Aβ, α-synuclein and tau. Available structures result from *in vitro* grown fibrils, *in vitro* amplified *ex-vivo* fibrils or directly from sarkosyl-extracted amyloids. The peptide segments that directly drive the amyloid assembly of these proteins have been previously identified and their structure solved mainly by X-ray diffraction of (micro)crystals of these peptides^11,14^. All α-synuclein polymorphs contain four such APRs (**Supplementary Fig. S1**). Aβ polymorphs invariably contain two APRs but here the amyloid propensity of the C-terminal APR is dependent on Aβ length (**Supplementary Fig. S2**). Tau also possesses two APRs although here both are not necessarily present together due to tau isoform variation (**Fig. 3A**). In all three amyloids, however, the stability of all polymorphs is dominated by their APRs forming stable quaternary ladders along the fibril axis which, as discussed above, is driven by their high cross-β propensity. This is also why they constitute the regions of highest local structural similarity between polymorphs. Polymorphism therefore results from the alternative packing of these APRs against each other and with less stable and non-amyloidogenic segments that are incorporated in these structures.

What is the reason for the ability of these proteins to form alternative stable tertiary interactions? First, the sequences of these proteins display relative low sequence entropy (i.e. sequence “richness”). The C-terminal APR of Aβ for example consists of 4 Gly, 3 Val, 3 Ile and 2 Ala while the low sequence entropy of tau and α-synuclein result from the fact that they consist of (imperfect) repeat sequences. As cross-β stacking produces arrays of perfectly aligned side chains this results in sterically very similar β-sheet surfaces. Second, both the structures of amyloidogenic peptide fragments, as well as full length fibrils show that most of the interaction energy between packing β-sheet interfaces can be attributed to 1 or two interdigitating (and often hydrophobic) side chain ridges. Together, these two features explain how APRs can both pack on themselves forming homozippers in small peptides but also establish heterotypic β-sheet packings in the full-length fibril. This also explains how non-amyloidogenic segments of the proteins are incorporated in the protofibrillar fold: packing of a single side chain ridge suffices to stabilize sequence segments that otherwise do not favour the cross-β conformation. This extra stabilisation is also apparent from the extension of the cross-β main-chain H-bond network beyond the boundaries of APRs although the H-bond geometry remains imperfect and therefore only partially saturated (**Fig. 3**).

Total surface contact between β-sheet elements is relatively similar in different polymorphs suggesting that their overall tertiary conformational stabilities are not very different. This implies that additional context-dependent interactions could easily steer polymorphic bias. Depending on buffer conditions, factors contributing to polymorphic bias of protofibrillar tertiary folds can include polymeric prosthetic ligands including proteoglycans, polyphosphates including RNA or DNA, lipid arrays, post-translational modifications or alternative modes of protofibrillar assembly into amyloid fibrils.

### In register side chain stacking favours amyloid specific metal and water binding sites

In order to obtain a more detailed analysis of the atomic interactions occurring in fibrillar structures compared to those observed in globular protein structures, we recorded distance-distribution profiles for all pairs of non-covalently bonded atoms in our collection of fibril structures and in the SCOP40 collection of globular structures (**Supplementary Table 1, sheet ‘Globular structures X-ray’**). We then computed the difference between each pairwise distance-distribution in fibrillar and globular structures and estimated the significance of the difference using a Kolmogorov-Smirnov test. To facilitate easy observation of the most significant differences, pairwise distance-distributions between fibrillar and globular structures were summarised in a volcano-style plot, with the difference between the interatomic distributions on the x-axis and the negative logarithm of the p-value on the y-axis (**Fig. 4A**). This analysis showed that the vast majority of interactions are similar between both classes of structures. As a first example, we show here the distributions between the backbone nitrogen and backbone oxygen atoms (**Fig. 4B**), i.e. hydrogen bond interactions as they typically occur in secondary structures, with the well-known peak at around 3Å distance. Of note, the fibrillar distance distribution is more focussed into a single peak, likely resulting from the single secondary structure (parallel β-sheet) occurring in these structures, whereas there is more diversity in the globular structures. As a second example, we show the packing between aliphatic carbons and aromatic nitrogen atoms, occurring at slightly higher than the sum of their van der Waals radii (**Fig. 4C**). However, the plot also revealed several pairwise distance distributions that differ significantly between globular and fibrillar structures, some interactions occur more frequently in fibrillar structures, while others occur much less frequently. For example, of the latter, particular backbone-nitrogen-to-sidechain-oxygen hydrogen bonds occur hardly at all in amyloids (**Fig. 4D and E**), likely because the backbone hydrogen bonds near-saturate the hydrogen bond potential. Other interactions that are depleted in amyloid structures are disulphide bonds and positive charge – aromatic pi-electron interactions (**Fig. 4A**). Conversely, we observe a notable reduction in amyloids of the occurrence of salt bridges between Arg and Lys side chains and their negatively charged Asp and Glu counterparts (**Fig. 4F and G**), especially since this is mirrored by a strong increase in proximity between atoms of similar charge, namely the positively charged nitrogens in Lys and Arg (**Fig. 4H**) or between the negatively charged oxygens in Glu and Asp (**Fig. 4I**). These distances indicate a stacking of same-charged residues, which derive from their superpositional stacking in the protofibril structures, where they form charged ladders that run along the length of the fibril (**Fig. 4F and G**). Given the strong charge repulsion that should be associated with this, it is unlikely that this source of structural frustration in these structures is easily compensated by the other interactions. It would appear more likely that either counterions lead to charge neutralisation or the pKa values of these carboxyl, amine and guanidine groups involved are strongly shifted to the same effect. Since neither the cryoEM nor the ssNMR structures contain direct information of the waters and ions, it remains unclear what is precisely going on in these structures with regard to these charge interactions. Of note, in many cryoEM structures, substantial additional densities have been noted that appear not to originate from the polypeptide, but rather from some associated other type of molecule^16,17^, suggesting that our current view on these amyloid structures is incomplete. The missing density is too large to be purely originating from solvent molecules and is likely to be one of several other classes of molecules that are known to influence amyloid aggregation *in vitro* and that have been found to co-deposit *in vivo*. These include lipids, metal ions and organic poly-ions, such as polyphosphates, polyamines and polysaccharides such as heparin. In particular, (poly)ionic molecules are of interest because of their strong association to amyloid deposition and the known effect of amyloids on metal homeostasis^18^.

**Figure 4.**
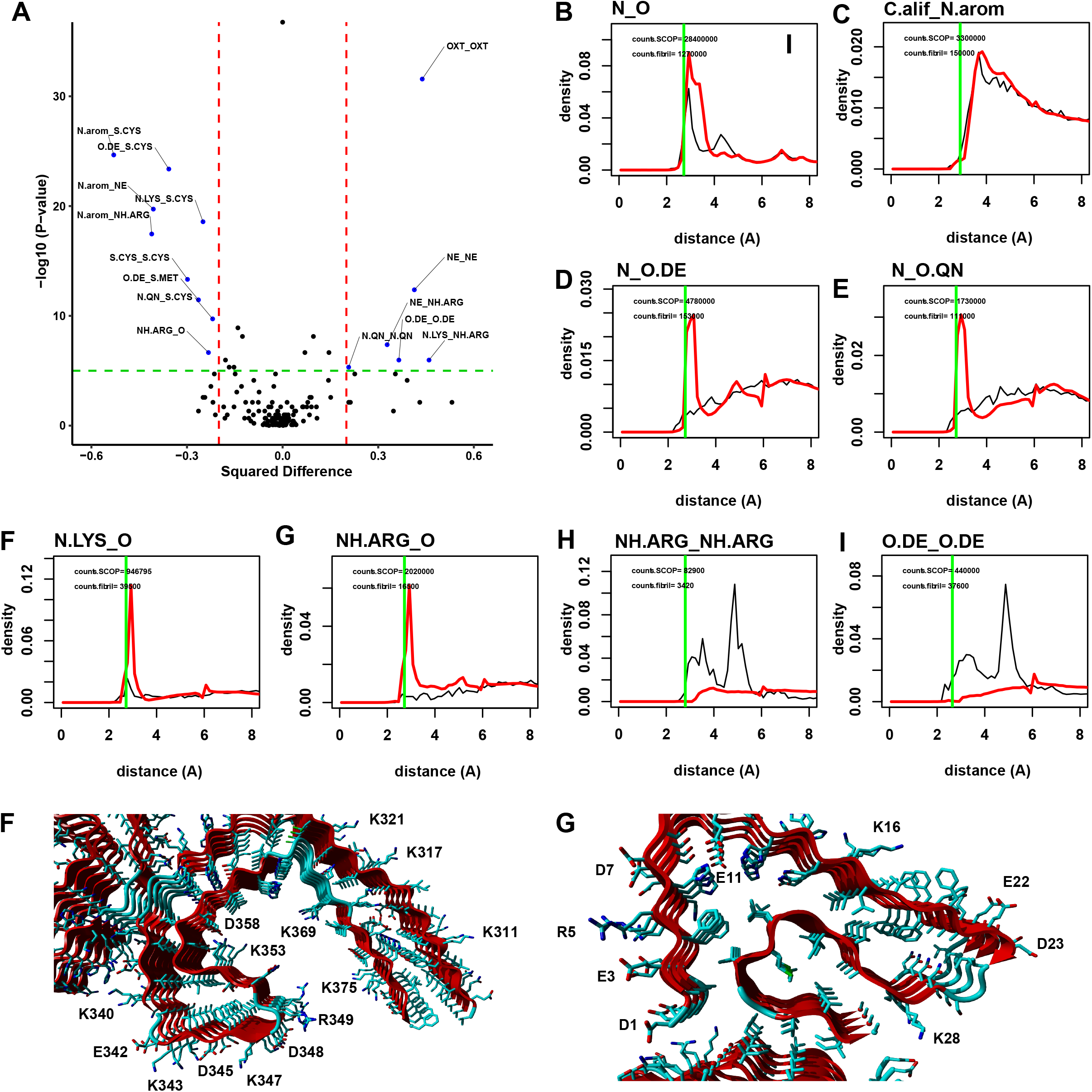
Profiling of atomic-atomic interactions in amyloid fibril structures. (**A**) Volcano plot highlighting interatomic distance comparisons between amyloid fibrils and globular proteins. Upper-right corner indicates enrichment of interatomic distances in amyloid structures, whereas the left-corner shows depletion. (**B-I**) Representative interatomic distance distributions that are similar (**B-C**), depleted (**D-G**) or enriched (**H-I**) in amyloid fibrils (shown in black lines) compared to globular folds (shown in red lines). Vertical green lines indicate the expected sum of VdW radii for each pair of atoms. (**F-G**). Examples of multiple charge ladders running along the axis of (**F**) tau (**PDB ID: 5O3L**) and (**G**) Aβ amyloid fibrils (**PDB ID: 5OQV**).

In particular for the solvent and metal ions, an algorithm is available in FoldX that predicts binding sites for structural waters and the most common metal ions in globular structures with good accuracy ^19^. The method is based on identifying positions on the protein surface where a water molecule or metal ion can form hydrogen bonds or electrostatic interactions with good geometry to multiple protein atoms. The method has been shown to predict more than 90% of the metal binding sites in globular proteins to within 0.6 Å, and 75% of the water molecules to within 0.8 Å ^19^. Utilising FoldX, we located multiple water and metal binding sites in the vast majority of these structures (**Fig. 5**). The number of tightly bound water molecules predicted per fibril structure ranges from 38 to 584, with a median of 204, forming ladders of structural waters running along the surface of the fibril. We found 1 water binding site per 1.7 residues in the fibrillar structures (56.682 waters predicted in 269 structures containing 96.645 amino acids in the fibril set), where this number of 1 per 0.96 residues in globular structures (416.314 waters predicted in 1911 structures containing 398.247 amino acids in the WHATIF set). So, although the set-size imbalance precludes a firm conclusion, it appears that fibrils binds less solvent than globular structures, likely reflecting their lower solubility. A typical solvent binding site in the fibrillar structures is composed of atoms from two consecutive layers of the fibril stack (**Fig 5A & 5B**), so that the water molecules act as bridges between the layers, likely stabilising the stacking further, much like structural waters stabilise globular protein structures. Similar to globular structures, both the backbone and side chains are involved in the coordination of water (**Fig 5C**).

**Figure 5.**
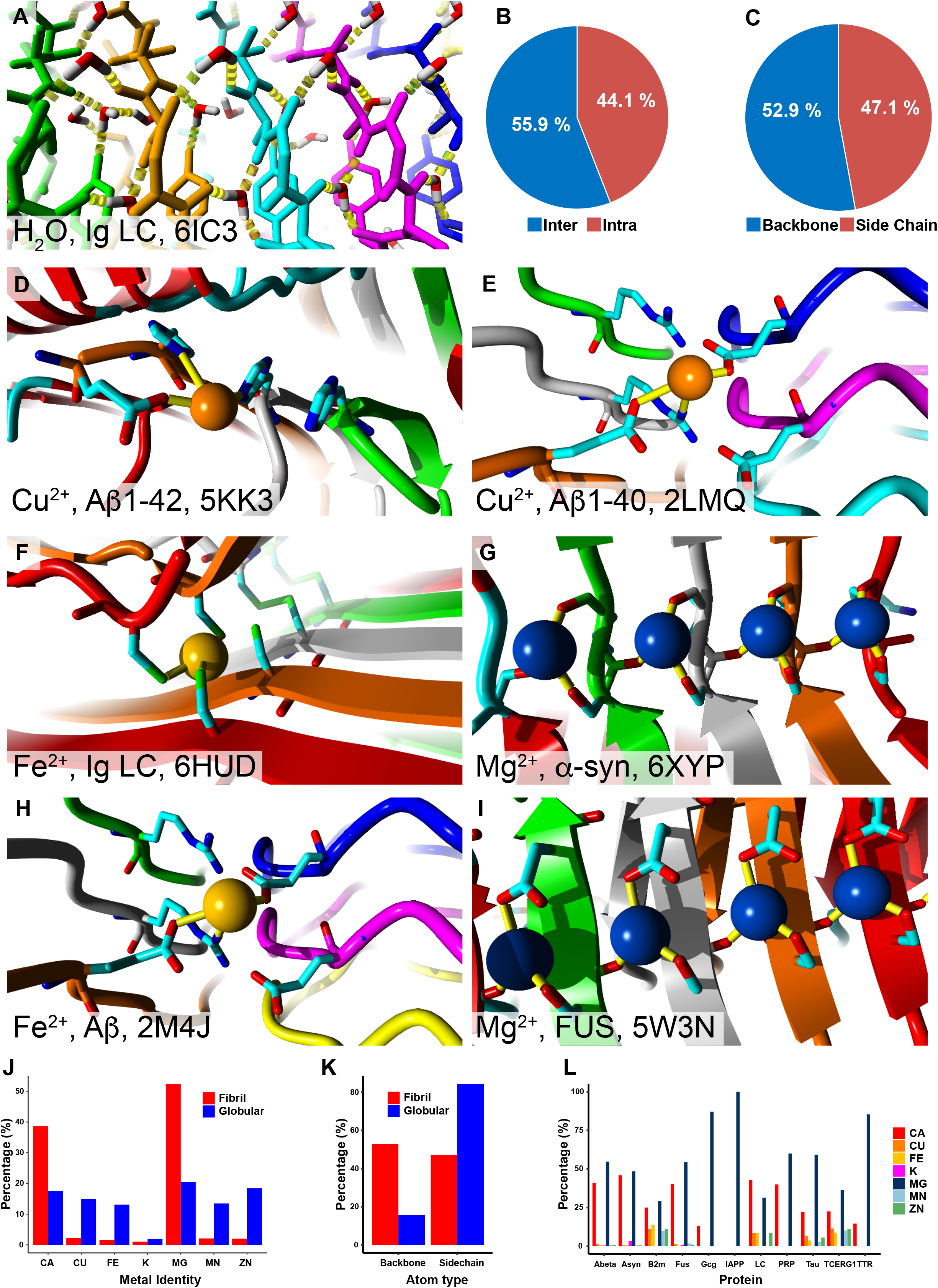
Predicted water and metal binding sites stabilise amyloid fibril structures. (**A**) Structural representation of multiple water binding sites in an amyloid-forming protein involving co-ordinating residues from multiple rungs. (**B**) Percentage of predicted waters coordinated by atoms from the same rung (Intra) compared to waters coordinated by atoms from multiple rungs (Inter). (**C**) Percentage of atoms belonging to the backbone or side chains interacting with water. (**D-I**) Structural representations of multiple metal binding sites in different amyloid-forming proteins involving co-ordinating residues from multiple rungs. (**J**) Amyloid fibril structures primarily bind to lower affinity metals Mg^2+^ and Ca^2+^ through (**K**) backbone atoms, compared to globular proteins. (**L**) Distribution of metal binding for different amyloid-forming proteins.

For the metals, the algorithm predicts binding sites in 133 out of 269 of the structures (**Fig 5D-5I**), with a strong bias towards the lower affinity metals Mg^2+^ and Ca^2+^, but others are also found at specific sites, including Cu, Fe, K, Mn and Zn (**Fig. 5J & 5K**). Similar to the solvent, there appear to be less metal coordination sites in fibrillar structures (1 metal ion predicted per 99 residues) than in globular structures (1 metal predicted per 52 residues). As with the solvent, we see most metal binding sites spanning 2 layers of the fibril stack, effectively interlinking the layers, thus likely providing further stabilisation of the amyloid structure. Moreover, there is a strong bias towards the involvement of backbone over side chain atoms in coordinating metal binding in fibrillar structures when compared to globular structures (**Fig. 5L**) and there are on average more atoms coordinating (almost always 4 residues). Taken together, a picture emerges of low affinity metal ion binding sites occurring as an intrinsic feature of the amyloid backbone arrangement, which is in contrast to the evolved high affinity metal binding sites in globular protein structures, which are mostly coordinated by side chains. From these analyses, it appears that both the solvent and the metal ions can be considered to be structural or intrinsic components of the amyloid fibrils, which explains the well-documented interplay between metal ion concentration and amyloid formation.

Although we do not have a good docking algorithm for the larger molecules that could bind to the amyloid fibrils, the regularity of the metal ion binding sites certainly supports the notion of poly-ionic polymers binding laterally across the layers of the fibril. This would explain the strong effect on amyloid kinetics of polymers, such as heparin or polyphosphate.

## Discussion

We have performed an analysis of over 60 amyloid structures. We find that the stability of amyloids is dominated by few short sequences displaying a high cross-β structural propensity representing about 30% of the residues in the amyloid core. These segments favour the in-register stacking of identical residues resulting in fully saturated main chain quaternary H-bond networks along the fibre axis. Remaining residues are either not contributing or even opposed to amyloid stability due to their lack of cross-β propensity. As a result, they either adopt unfavourable dihedral angles that lack main chain H-bond saturation or adopt forced β-sheet conformation which is apparent from suboptimal main chain H-bond geometries. Unfavourable sequence segments are nevertheless incorporated in the proteolysis resistant core of amyloids due to additional stabilizing interactions. This is often achieved by packing of unstable β-sheet interfaces against the sheet of more stable segments. We find that in such cases most of the interaction energy results from the interdigitation of only one or two side chain ridges of both β-sheet surfaces. This focal stability of tertiary β-sheet packing interactions in few intercalating ridges is one of the important reasons for the existence of polymorphism. Indeed, combined with low sequence entropy and a repeat sequence structure this explains why the tertiary structure of Aβ, α-synuclein and tau can adopt such a variety of alternative packings. Given that the total buried surface in different tertiary packings is relatively similar, we can assume that alternative tertiary polymorphs have relatively equivalent thermodynamic stabilities. This suggests that without additional directive interactions, polymorphism can be expected to be degenerate. Polymorphic ambiguity is exactly what is observed for in vitro grown recombinant tau^20,21^. Recombinant tau contains both APRs and can adopt a variety of polymorphs in simple aqueous buffers *in vitro* including the 4R-twister, 4R-snake and 4R-jagged conformations^21^. Interestingly, here polymorphic ambiguity is not only observed between different fibrils in the same sample but polymorphic transitions between the different polymorphs can also be observed within single fibrils. Additionally, it is also becoming clear that polymorphism is not necessarily conserved when growing *in vitro* fibrils from patients, e.g. for α-syn from MSA patient^22^. Our analysis shows how polymorphs share a similar stability profile along the fibril axis that is determined by a common framework of stable cross-β forming segments, while displaying ambiguous yet energetically equivalent tertiary folds across the fibril axis. It can therefore be envisaged that intrafibrillar polymorphic transitions can result from conservation of the APR framework, which is also required for fibril elongation but can be accompanied by a reconfiguration of more unstable regions. The same mechanism could also explain the occurrence of fibrillar imperfections that are favoured by conditions of fast elongation and are believed to constitute fibril fragmentation sites or sites of secondary nucleation occurring at the rate of 1:100 to 1:150 rungs under typical *in vitro* conditions of fibril elongation^23^. Which factors then contribute to polymorphic bias and enhanced polymorphic conservation observed in disease? First, the same spectrum of available amyloid conformations can be limited by sequence isoforms: *in vivo* tau displays 3R,4R and mixed 3R/4R polymorphs and Aβ peptides come in lengths varying from 36 to 46 amino acids^24,25^. Second, post-translational modifications both at positions inside the amyloid core but also outside e.g. in the fuzzy coat will favour specific polymorphs^24^. Third, most patient extracted amyloid structures contain non-proteinaceous ligands that often dock close to unstable segments of the fibrils thereby favouring a specific tertiary conformation^26^. Although most prosthetic ligands are still unidentified, potential candidates include polyionic molecules including various glycosaminoglycans, polyphosphates including RNA, arrayed lipids and others^27,28^. Thus, although in reality not all amyloids will have the same polymorphic instability as observed for *in vitro* grown recombinant tau^21^, it is still probable that polymorphic bias is partially (re)enforced by specific cellular and physiological contexts. Our thermodynamic analysis suggests that the specific physiological or pathological interactions in vulnerable cells could facilitate the nucleation and elongation of specific polymorphic conformations, first possibly by capturing more fluid soluble oligomers. The polymorphic bias introduced in this manner would imply that once formed seeds would be entropically primed to interact more efficiently with similar cellular components after propagation to successive vulnerable cells. It also implies that, in order to maintain polymorphic bias, these interactions need to be present in successive cells (**Fig. 6**). Together this could explain the disease-specific tropism of different polymorphs^11,14,16,20-22,29-31^. Of course, amyloid plasticity also implies that amyloid structure is likely affected by solvents and detergents used for tissue extraction and it therefore remains unclear which are the relevant structures in disease. Nevertheless, the thermodynamic principles apparent from the amyloid structures and their polymorphs analysed here are likely conserved.

**Figure 6.**
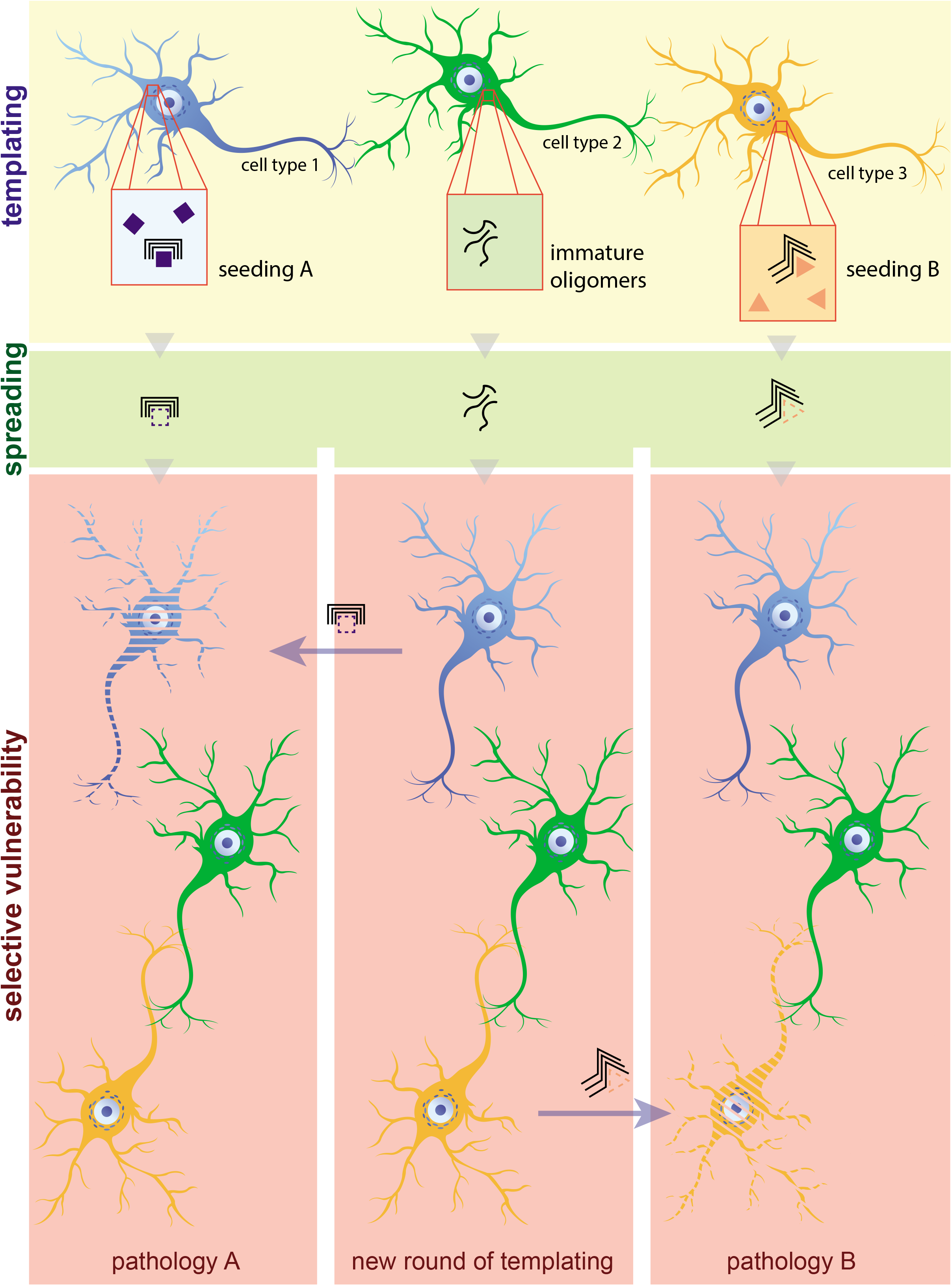
Polymorphic bias as a mechanism promoting cellular susceptibility. The templating bias that structural polymorphs exhibit explains how certain cellular types sharing similar intrinsic polymorphic components are particularly vulnerable to amyloid spreading of entropically primed seeds (left and right column) and why particular polymorphs are tethered to certain forms of disease (seeding A to pathology A, seeding B to pathology B). This also precludes that the polymorphic content of cells may also potentially capture and shape unspecific early oligomeric forms to promote an additional layer of biased spreading that further enhances cellular vulnerability and distinct pathologies (middle column) at a second level.

Although we cannot currently identify the exact nature of amyloid ligands in known structures, FoldX allows us to model both metal binding sites and structural waters in amyloid structures. We found that amyloids contain a significant amount of metal and water binding sites, particularly at partially or non-satisfied main-chain H-bond positions between rungs, thereby additionally stabilizing the amyloid fibrils. Due to the repetitive nature of fibrils, this results in a high local concentration of potential metal binding sites and it is therefore plausible that in particular instances metal avidity during the process of amyloid elongation results in the local dysregulation of metal homeostasis^32,33^. We also find that although amyloids bind less molecules of water than globular proteins per polypeptide chain unit, like metals they do form arrayed lattices of structural waters. This might explain why amyloids often display antifreeze properties^34^. It also suggests that amyloids and globular proteins will have very different water chemical potentials and will be very differently affected by osmolytes.

Finally, the thermodynamically stable segments identified here largely overlap with aggregation-prone segments (APRs) previously found to be necessary for amyloid nucleation and elongation^10,11,14^. This does not exclude modulating effects of other regions, even by regions that are not incorporated in the fibre core, such as the fuzzy coat observed in many amyloids^29^. However, by and large in disease amyloids assembly kinetics and stability are both determined by the same aggregation-prone regions. This is radically different in globular proteins where regions determining protein stability and folding kinetics are disjointed. In globular proteins, interactions forming the folding nucleus are established first while stabilizing interactions are established only after the rate-limiting step of folding. This could have important implications for the evolution of globular protein folds. Indeed, many amyloids populate soluble oligomeric conformations, some of which adopt globular-like structure. Our analysis shows that the propensity of disease amyloids to form soluble oligomers is indeed probable given the high degree of structural frustration observed in amyloids. The emergence of stable globular structure from amyloidogenic peptide sequences would therefore only require a limited degree of kinetic adaptation. Mutations slowing amyloid assembly (e.g. gatekeeper mutations at the flanks of APRs) together with mutations favouring more rapid alternative modes of assemblies (e.g. out of register β-sheet assembly) would be sufficient to drive that process^35-37^.

## Online Methods

### Collection of amyloid fibril structures

We collected a complete set of recently published amyloid fibrils structures solved by ssNMR or cryoEM from the PDB^38^. The structures include, among others, several polymorphs of the Alzheimer β-peptide 1-40^39-44^ (Aβ40) and 1-42^45-50^ (Aβ42), alpha-synuclein^17,26,51-57^ (α-syn) and tau^16,20,21,29-31^, as well as single polymorphs of β-2-microglobulin^58,59^ (β2M), Islet Amyloid Polypeptide^60^ (IAPP), and Serum Amyloid A1^61^ (SAA1) (**Fig. 1 and Supplementary Table 1**).

### Quality assessment of structural datasets

To gain more insight into the determinants of amyloid fibril architecture, we analysed the thermodynamic profile of 66 available amyloid structures (**Supplementary Table 1, sheet ‘Fibril structures’**) and compared them to a set of 1926 high quality globular protein structures, selected from the PDB, using Whatif^62^ (R-factor<0.19 and Resolution<1.5 Å, and 90% sequence homology (**Supplementary Table 1, sheet ‘Globular structures X-ray’**). In order to perform this analysis, we used the FoldX force field^6^. The method was described previously ^63,64^, but briefly, during a free energy calculation, FoldX first calculates the free energy contribution of each atom in the protein based on its position relative to its neighbours in the structure. These contributions are then summed, first at the residue level and later to the level of the entire protein. This allows to map the contribution of each residue to the total free energy of the protein (called G_contrib_) but also to report on individual thermodynamic components (e.g. Van der Waals interactions, electrostatics, H-bonding or electrostatics, enthalpy) contribute to the stability of the structure.

The accuracy of the free energy estimation by FoldX correlates with structure quality parameters, such as crystallographic resolution. The quality of protein polypeptide backbone conformations can easily be estimated by plotting the Ramachandran map of these structures and evaluating whether backbone traces deviate from the allowed conformational space of each of its residues. In order to do this in a quantitative manner, FoldX uses a statistical thermodynamics approach based on the observed frequency distribution of each amino acid in the Ramachandran map in crystal structures of the highest resolution (selected using the WHATIF tool^62^). Given the overall lower resolution of cryoEM structures in comparison to high quality crystal structures this will still result in a lower accuracy of energetic calculations by FoldX. In addition, cryo-EM structures often display undetermined non-proteinous regions of structural density which will not be considered by FoldX. However, and despite these caveats, given the overall good quality of cryoEM backbone conformations the accuracy of energetic calculations can be expected to be comparable to the quality achieved for low sequence identity (40-50%) homology modelling which is sufficient to correctly identify the residues and/or structural regions of a protein that are key to its thermodynamic stability.

### Energy profiling of amyloid structures

To analyse the regional stability of amyloid structures, we used FoldX to calculate the contributions of individual residues to the thermodynamic stability of the structure (ΔG_contrib_) and plotted these energy profiles along the sequence for each structure. In order to smoothen the energy profile and allow correlation with physical properties, we averaged the values over a sliding window of size 5 and linearly rescaled the values to a mean of 0 and standard deviation of 1. For comparison, we include all energy profiles for all polymorphs of a protein in a single plot (**Fig. 3A and Supplementary Fig. S1 & S2**).

### Receiver-Operator curve (ROC) analysis

ROC curves were determined by plotting the fraction of correct predictions (True Positive Rate) against the fraction of wrong predictions (False Positive Rate) for all the per residue values of the predicting variable (hydrophobicity or β-sheet propensity in this case). From such plots, the area under the curve (AUC) characterises the overall performance of the predictions. As a second measure, using the ROC curves we also calculated the optimal Matthews Correlation Coefficient for both structural features. The R package ROCR was utilising to calculate and plot the corresponding ROC curves^65^.

## Acknowledgements

This work was supported by the Flanders institute for Biotechnology (VIB); KU Leuven; the Fund for Scientific Research Flanders (FWO, project grants G0C2818N, G0C0320N and G053420N); the Stichting Alzheimer Onderzoek (SAO-FRA 2019/0015, SAO-FRA 2020/0009 and SAO-FRA 2020/0013); the European Research Council under the European Union’s Horizon 2020 Framework Programme ERC Grant Agreement (647458 MANGO); NL was also funded by Fund for Scientific Research Flanders Postdoctoral Fellowship (12P0919N).

