## Supplementary Figures 1 and 2 and legends for "A structural analysis of amyloid polymorphism in disease: clues for selective vulnerability?"

van der Kant & Louros, Supplementary Figure 1

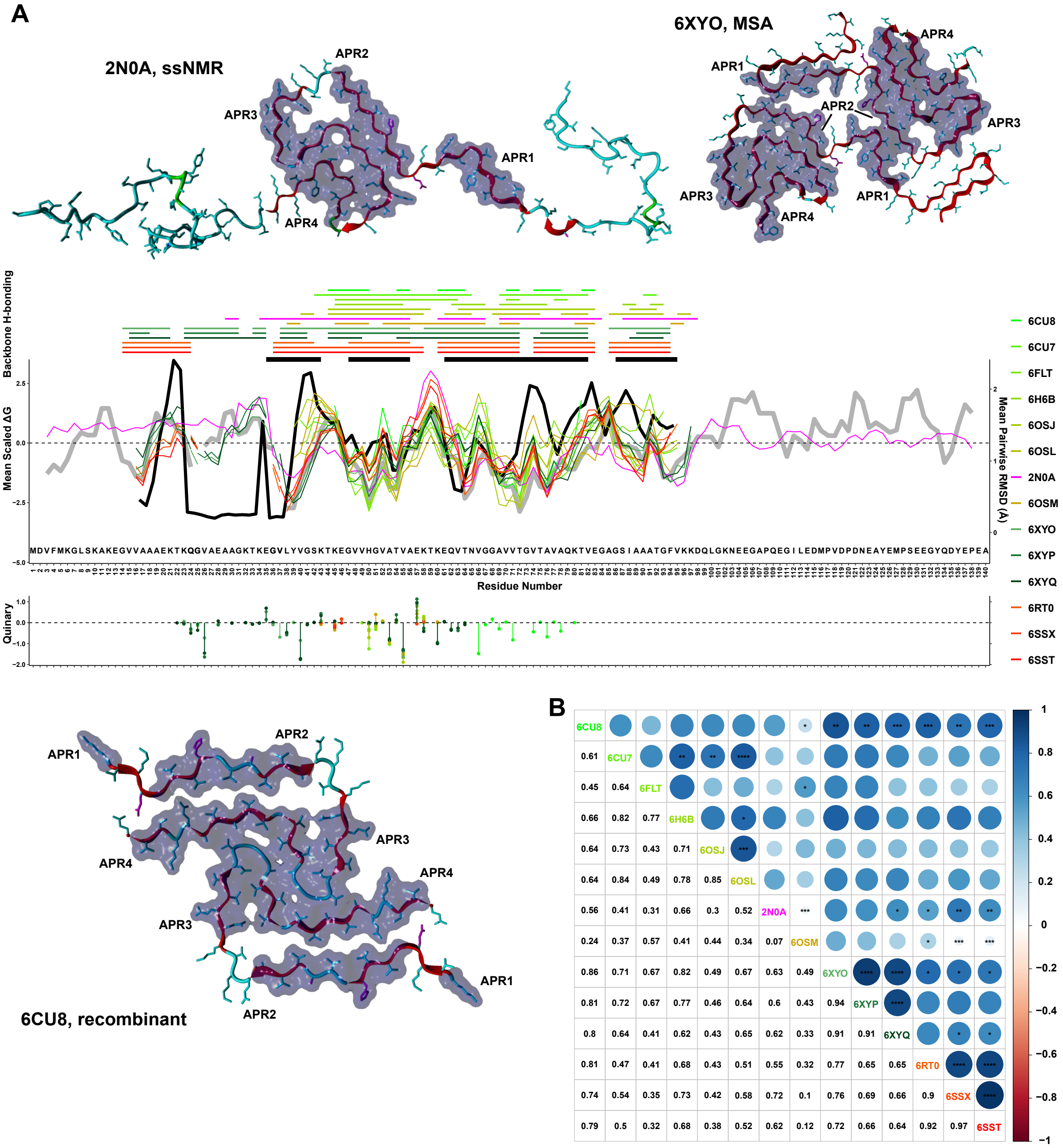


**Figure S1. Cross-comparison of defining features between amyloid structural polymorphs of α-syn.** (**A**) Representative structures of α-syn polymporhs, with positions of known APR segments highlighted as black segments. Distributions of stability free energy contributions (middle plot) are aligned to quinary energy contributions (bottom plot) and overlaid in different colours for different structural polymorphs of α-syn. The top plot indicates hydrogen bond patterning (colour-coded per polymorph) and the position of experimentally determined APRs (black segments). RMSD distribution, produced after superimposing all structures, is plotted as a thick black line in the middle plot. (**B**) Pairwise comparison plot of individual α-syn polymorphs. Top-half indicates strong structural similarities by circle-sizes (colour-coded from red to blue) and bottom half indicates pairwise scores.

van der Kant & Louros, Supplementary Figure 2

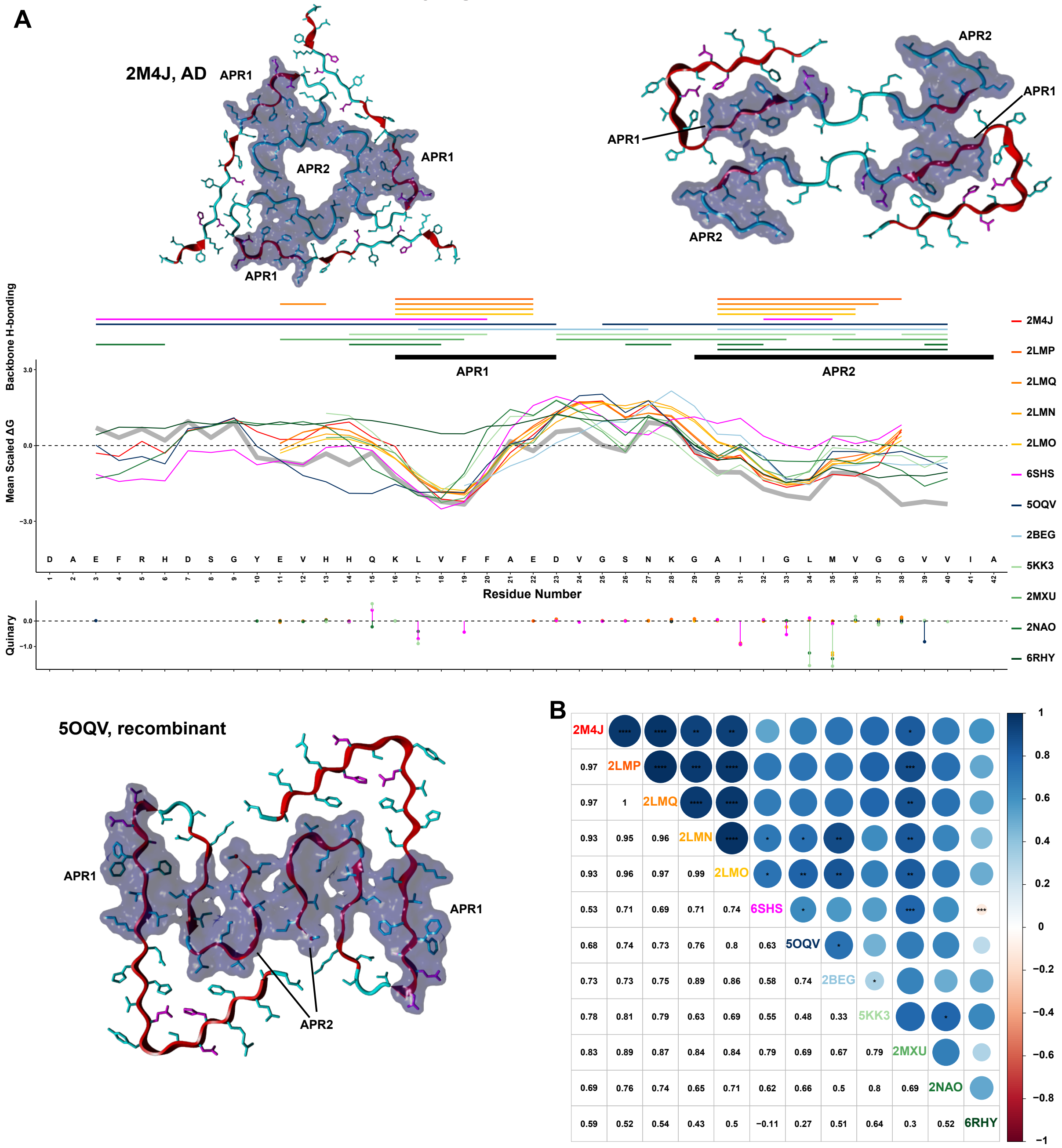


**Figure S2. Cross-comparison of defining features between amyloid structural polymorphs of Aβ.** (**A**) Representative structures of Aβ polymporhs, with positions of known APR segments highlighted as black segments. Distributions of stability free energy contributions (middle plot) are aligned to quinary energy contributions (bottom plot) and overlaid in different colours for different structural polymorphs of Aβ. The top plot indicates hydrogen bond patterning (colour-coded per polymorph) and the position of experimentally determined APRs (black segments). (**B**) Pairwise comparison plot of individual Aβ polymorphs. Top-half indicates strong structural similarities by circle-sizes (colour-coded from red to blue) and bottom half indicates pairwise scores.
